# Do bed bugs transmit human viruses, or do humans transmit bed bug viruses? A worldwide survey of the bed bug RNA virosphere

**DOI:** 10.1101/2023.10.20.563367

**Authors:** Hunter K. Walt, Jonas G. King, Johnathan M. Sheele, Florencia Meyer, Jose E. Pietri, Federico G. Hoffmann

## Abstract

Bed bugs (Hemiptera: *Cimicidae*) are a globally distributed hematophagous pest that routinely feed on humans. Unlike many blood-sucking arthropods, they have never been linked to disease transmission in a natural setting, and despite interest in their role as disease vectors, little is known about the viruses that bed bugs naturally harbor. Here, we present a global-scale survey of the bed bug RNA virosphere. We sequenced the metatranscriptomes of 22 individual bed bugs (*Cimex lectularius* and *Cimex hemipterus*) from 8 locations around the world. We detected sequences from two known bed bug viruses (Shuangao bedbug virus 1 and Shuangao bedbug virus 2) which extends their geographical range and the host range of Shuangao bedbug virus 1 to *Cimex lectularius*. We identified three novel bed bug virus sequences from a tenui-like virus (*Bunyavirales*), a toti-like virus (*Ghabrivirales*), and a luteo-like virus (*Tolivirales*).

Interestingly, some of the bed bug viruses branch near to insect-transmitted plant-infecting viruses, opening questions regarding the evolution of plant virus infection. When we analyzed the putative viral sequences by their host’s collection location, we found unexpected patterns of geographical diversity that may reflect humans’ role in bed bug dispersal. Additionally, we investigated the effect that *Wolbachia,* the primary bed bug endosymbiont, may have on viral abundance and found that *Wolbachia* infection neither promotes nor inhibits viral infection. Finally, our results provide no evidence that bed bugs transmit any known human pathogenic viruses.

## 1. INTRODUCTION

Bed bugs (Hemiptera: Cimicidae) are globally distributed obligately hematophagous ectoparasites (Doggett et al., 2012). Some species routinely feed on humans, but unlike many other blood sucking insects, there is no evidence that bed bugs are human disease vectors (Davies et al., 2012). Several studies have been conducted to assess if bed bugs could transmit human pathogens, but transmission in the wild has not been documented (Blakely et al., 2018; El Hamzaoui et al., 2019; Goddard & deShazo, 2009; Leulmi et al., 2015; Pietri, 2020). Considering the consistent resurgence of bed bug populations leading to outbreaks in the last 20-30 years, it is important to understand what microbes they harbor along with their potential as vectors (Davies et al., 2012; Doggett & Lee, 2023; Lewis et al., 2023).

Large scale metatranscriptomic studies have enhanced our knowledge of viral diversity in invertebrates and have made detecting arthropod-associated viruses increasingly more feasible (Käfer et al., 2019; Li et al., 2015; Shi et al., 2016). Bed bugs have been included in some arthropod viromics-based studies, and four putative bed bug virus sequences have been detected: Serbia reo-like virus 2 (NCBI:txid2771464), Serbia picorna-like virus 2 (NCBI:txid2771462), Shuangao bedbug virus 1 (NCBI:txid1608071) and Shuangao bedbug virus 2 (NCBI:txid1608072) (Li et al., 2015; Zhang et al., 2020). No further studies have been conducted to examine the pathogenic properties of these putative viruses, but they do provide evidence that bed bugs encounter viral infection. Furthermore, Ling *et al*. (2020) took a metatranscriptomic approach to survey for viruses in bed bugs and found that some reads in a sample of recently blood fed individuals aligned to hepatitis C virus. Although there was no evidence that the virus was replicating (i.e., low number of reads mapped and an incomplete genome), their study highlights the role that bioinformatic surveillance could play in detecting human pathogen transmission in bed bugs. However, bed bug specific viruses were either not detected or not reported in their study (Ling et al., 2020).

In this study, we conducted a worldwide survey of the bed bug RNA virosphere. Our aims were to search for known human viral pathogens and novel bed bug viruses that could be of interest to human health or biocontrol. We collected bed bugs (*Cimex lectularius* and *Cimex hemipterus*) from 8 distinct locations around the world and sequenced RNA libraries from 22 individuals. We assembled bed bug metatranscriptomes, and conducted phylogenetic analyses on the virus sequences that we detected. We also assessed viral diversity between bed bug species and geographic location. Additionally, we investigated whether there is a correlation between *Wolbachia* (a bed bug endosymbiont known to have an antiviral effect when infecting other insect taxa) reads and bed bug virus reads in each sample (Cogni et al., 2021; Hussain et al., 2023; Lindsey et al., 2018; Teixeira et al., 2008; Terradas & McGraw, 2017).

## 2. MATERIALS AND METHODS

### 2.1. Collection and extraction

The samples used in this study were obtained as part of a large international collection of bed bugs provided by numerous pest control companies and researches. We included *Cimex lectularius* samples from Czechia (n=3), France (n=1), the UK (n=3), Rome-Italy (n=3), Assisi-Italy (n=3), Ohio-USA (n=3), the Harlan lab strain of *Cimex lectularius* (initially collected in Fort Dix, New Jersey, USA and maintained at the King lab at Mississippi State University) (n=3), and *Cimex hemipterus* samples from Madagascar (n=3). Each bug was washed in 95% ethanol and total RNA was isolated using a standard TRIzol protocol (Invitrogen Waltham, MA) and further purified using NEB’s Monarch RNA cleanup kit was used (New England Biolabs Ipswich, MA).

### 2.2. Library Prep and Sequencing

We checked total RNA quality using Nanodrop quantitation and agarose gel electrophoresis, and we assessed RNA integrity with an Agilent 2100 bioanalyzer. Libraries were prepared using a strand specific library prep with ribosomal RNA depletion. Sequencing was conducted on the Illumina NovaSeq 6000 instrument for 150 base pair paired-end reads, resulting in approximately 100-135 million reads per sample. The quality of the reads was inspected using FastQC v0.11.5 (Andrews, 2010) and reads were quality trimmed using trimmomatic v. 0.39 (Bolger et al., 2014).

### 2.3. Virus Discovery

To enrich the data for viral reads, we filtered out *Cimex*, *Wolbachia*, and human reads by mapping to the *Cimex lectularius* (GCF_000648675.2), *Cimex hemipterus* (GCA_001663875.1, partial), *Wolbachia* endosymbiont of *Cimex lectularius* (GCF_000829315.1), and *Homo sapiens* (GCF_000001405.40) genomes using the bbsplit.sh tool of the bbmap suite (version 38.46) (sourceforge.net/projects/bbmap/) and retained all unmapped reads. We assembled the unmapped reads using Trinity v.2.11.0 (Grabherr et al., 2011) both individually and by collection location. We clustered the individual sample assemblies with cd-hit-est (W. Li & Godzik, 2006) with a sequence similarity threshold of 90%, a minimum sequence length of 500 nt, and a word size of 8. The assemblies were annotated using diamond blastx in the –very-sensitive mode with an e-value cutoff of 1e^−5^ (Buchfink et al., 2015, 2021) and the annotated transcripts were filtered for viral hits which were further inspected for false positives using NCBI’s BLASTx (https://blast.ncbi.nlm.nih.gov/Blast.cgi) against the nr protein database. To detect known viral conserved domains, we also inspected each viral hit in NCBI’s conserved domain database (CDD) (https://www.ncbi.nlm.nih.gov/Structure/cdd/cdd.shtml) and InterProScan (https://www.ebi.ac.uk/interpro/search/sequence/) (Quevillon et al., 2005).

When putative multipartite virus sequences were present, we used a viral co-occurrence detection method we previously described to identify the other genomic segments (Walt et al., 2023). Briefly, this method calculates two metrics based off sample co-occurrence. The first metric, *V_co_*, determines the frequency at which a transcript occurs in a sample with a given viral conserved sequence (e.g., transcripts with confirmed RdRp domains detected in BLAST analysis). The second metric, *T_co_*, determines if the transcripts found together with a viral conserved sequence occur in other samples without the viral conserved sequence. We used the thresholds *V_co_*=0.75 and *T_co_*=0.5 to determine candidate viral genomic segments. After running the co-occurrence analysis, we only kept candidate sequences with an ORF size > 500 nt. We further inspect these transcripts for conserved protein families and domains using NCBI’s CDD and InterProScan.

### 2.4 Phylogenetic Analysis of Viruses

We only used transcripts with confirmed RdRp domains for phylogenetic analyses. First, we extracted and translated ORFs encoding for RdRp proteins using NCBI’s ORFfinder (https://www.ncbi.nlm.nih.gov/orffinder/) and we downloaded diverse representative sequences from families within relevant viral orders from NCBI. We aligned RdRp amino acid sequences using MAFFT v7.490 (using the E-INS-I, G-INS-i, and L-INS-i, algorithms) and MUSCLE v3.8.1551 (Edgar, 2004; Katoh & Standley, 2013). We compared alignment qualities using MUMSA (Lassmann & Sonnhammer, 2006) scores, and the highest scoring alignment was used for tree inference. Phylogenetic analyses were conducted with IQ-TREE2 v.2.0.7 using ModelFinder (Kalyaanamoorthy et al., 2017) to find the best fitting substitution model. We assessed branch support using ultrafast bootstrap with 1000 replicates, Shimodaira-Hasegawa-like approximate likelihood ratio test (SH-aLRT) with 1000 replicates, and the aBayes test (Anisimova et al., 2011; Minh et al., 2013; Nguyen et al., 2015). All phylogenetic trees were midpoint rooted (unless otherwise noted) and visualized using the Interactive Tree of Life (iTOL) webserver (Letunic & Bork, 2021).

### 2.5 Phylogeographic Analysis

We retrieved all transcripts of putative viral origin from each individual bed bug assembly using Diamond BLASTx and CD-hit output. We predicted coding sequences using EMBOSS’s getorf tool using the -find 2 option. We only used coding sequences with complete RdRp domains for phylogenetic analysis and duplicate sequences within samples were discarded. We conducted phylogenetic analysis in the same way as section 2.4, except that nucleotide sequences were used instead of amino acid. We selected closely related taxa from the analysis in 2.4 as outgroups for phylogeographic analysis. We calculated evolutionary distance as p-distance using MEGA v.11.0.13 (Tamura et al., 2021). We did not conduct phylogeographic analysis on Shaungao Bedbug virus 2 because it was only detected in one location in our study.

### 2.5 Correlation of Wolbachia and Viral Read Abundance

We mapped the trimmed read datasets to all the viral genomes detected in this study and the *Wolbachia* endosymbiont of *Cimex lectularius* genome (GCF_000829315.1) using HISAT2 v.2.2.1 (Kim et al., 2019). We used the summary file output to obtain the percent of reads that mapped to the virus genomes or the *Wolbachia* genome. To correlate the Wolbachia and virus read abundances, we conducted a simple linear regression analysis in R v.4.2.2. (R Core Team, 2020), using the percentage of reads that mapped to the viral genomes versus the percentage of reads mapped to the *Wolbachia* genome.

## 3. RESULTS

### 3.1 Read Mapping, Assembly, and Transcript Annotation

To survey the bed bug virome, we sequenced the total RNA (ribosomal and small RNA depleted) from 22 individual bed bugs collected from the United Kingdom, France, Chechia, Italy, Madagascar, and the USA. To enrich for virus sequences in our bed bug samples, we filtered out reads that mapped to genomes of organisms that could be represented in our dataset. We classified an average of 85% of reads from each sample as bed bug (65.2% *C. lectularius*, 19.8% *C. hemipterus*), *Wolbachia* (2.08%), or human reads (0.07%). Those reads were not used for metatranscriptome assembly (**Figure 2**). We assembled the remaining reads and aligned the assemblies to NCBI’s nr protein database using diamond BLASTx, which returned multiple significant hits to putative bed bug virus sequences. We detected sequences from two known negative-sense single stranded RNA (-ssRNA) viruses associated with *C. hemipterus*: Shuangao bed bug virus 1 (Sbbv1) and Shuangao bed bug virus 2 (Sbbv2), but we did not detect the other two known bed bug viruses in our assemblies (C. X. Li et al., 2015; Zhang et al., 2020). In addition, we detected 3 novel viral sequences. These sequences all belong to the realm *Riboviria* and encode for 1.) a -ssRNA genome, 2.) a positive-sense single stranded RNA (+ssRNA) genome, and 3.) a double stranded RNA (dsRNA) genome. Strikingly, all putative viral sequences detected in this study are present intercontinentally, but none were detected across all samples.

**Figure 1:**
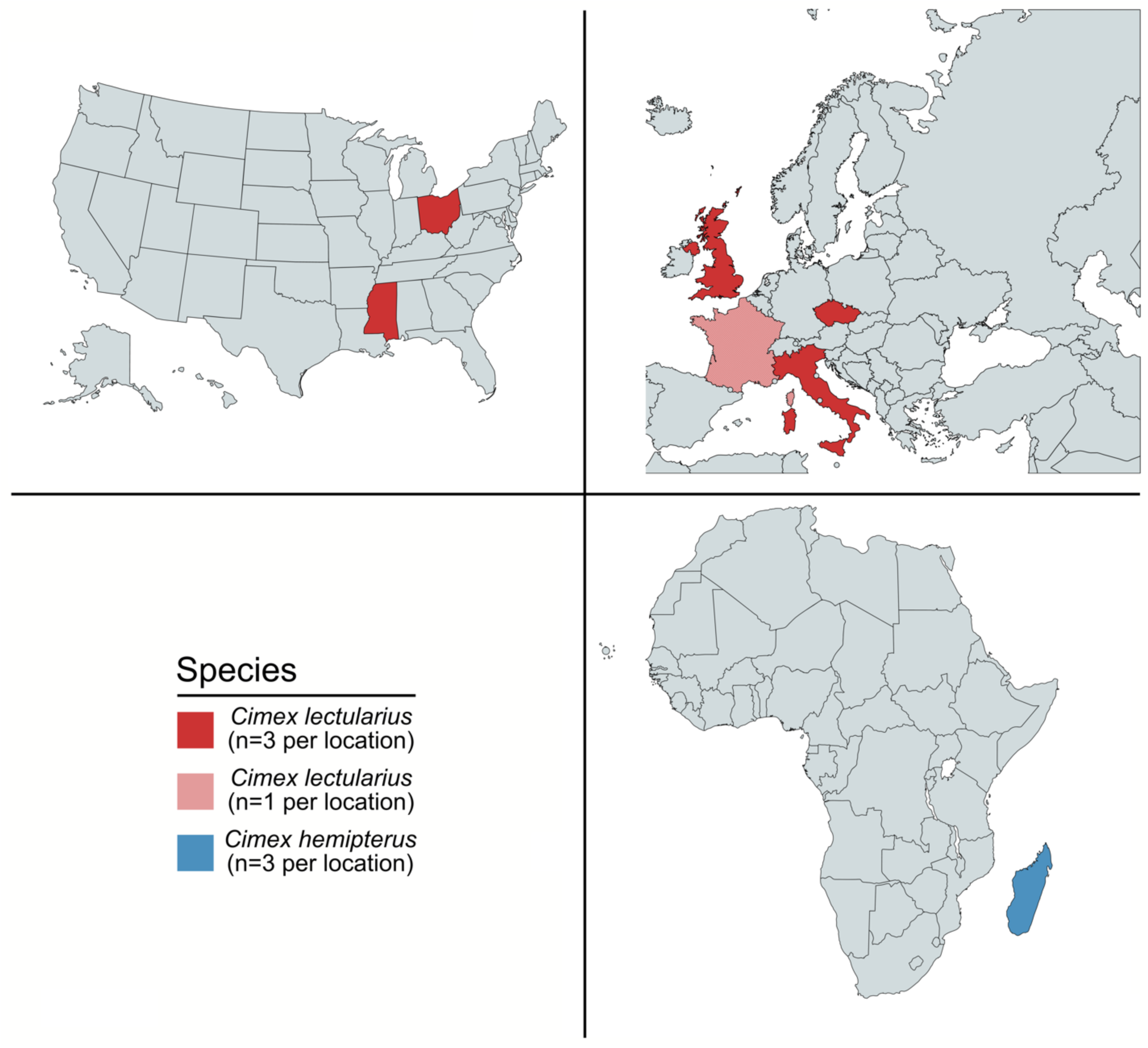
Locations of sample collection sites. Samples were collected at nine distinct sites around the world: France (*Cimex lectularius,* n=1), the UK (*Cimex lectularius,* n=3), Czechia (CR) (*Cimex lectularius,* n=3), Rome-Italy (*Cimex lectularius,* n=3), Assisi-Italy (*Cimex lectularius,* n=3), Madagascar (*Cimex hemipterus,* n=3), Ohio-USA (*Cimex lectularius*, n=3), and the King lab at Mississippi State University-USA (Harlan Strain-USA *Cimex lectularius,* n=3).

**Figure 2:**
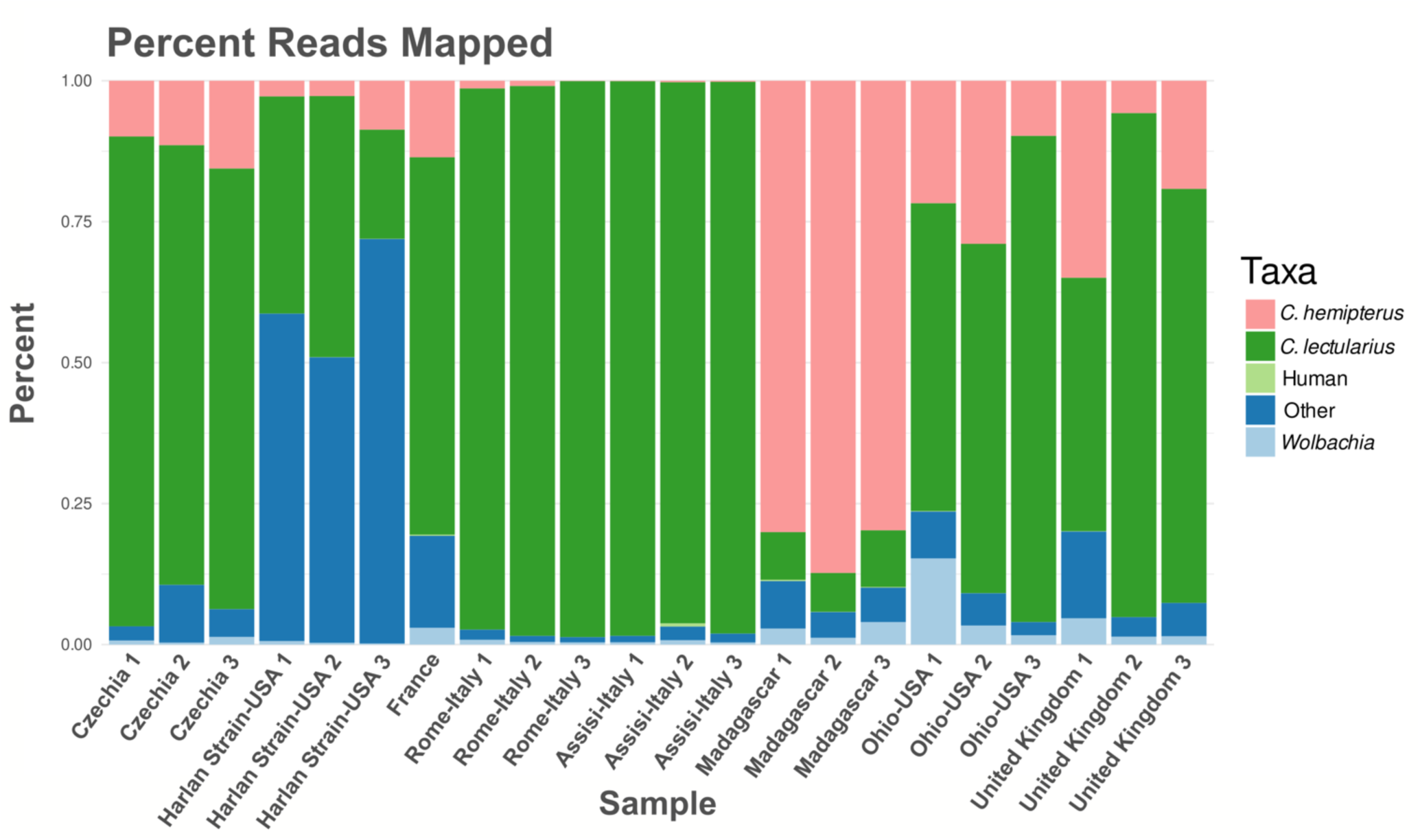
Percentage of reads mapped to the bed bug genomes, human genome, and *Wolbachia* genome per sample.

### 3.2 Shuangao Bedbug Virus 1

Sbbv1 is described as an unclassified rhabdovirus (-ssRNA) (NCBI:txid1608071) in the NCBI database. Our phylogenetic analysis groups Sbbv1 within the *Bunyavirales* order branching sister to the *Hantaviridae,* which is consistent with previous studies (C. X. Li et al., 2015) (**Figure 3**). We detected Sbbv1 sequences in *C. hemipterus* samples from Madagascar, and *C. lectularius* samples from France and Czechia. Our results extend Sbbv1’s geographical range from China (where it was first detected) to Madagascar, Czechia, and France, and its host range to *C. lectularius*. We detected both known segments of Sbbv1: a ∼7000 nt segment (L) containing the RdRp domain, and a ∼3000 nt (M) segment encoding for the glycoprotein precursor. The complete M segment was detected in Czechia sample 3, France, and all three Madagascar samples, but it was only partially complete in Czechia Sample 2.

**Figure 3:**
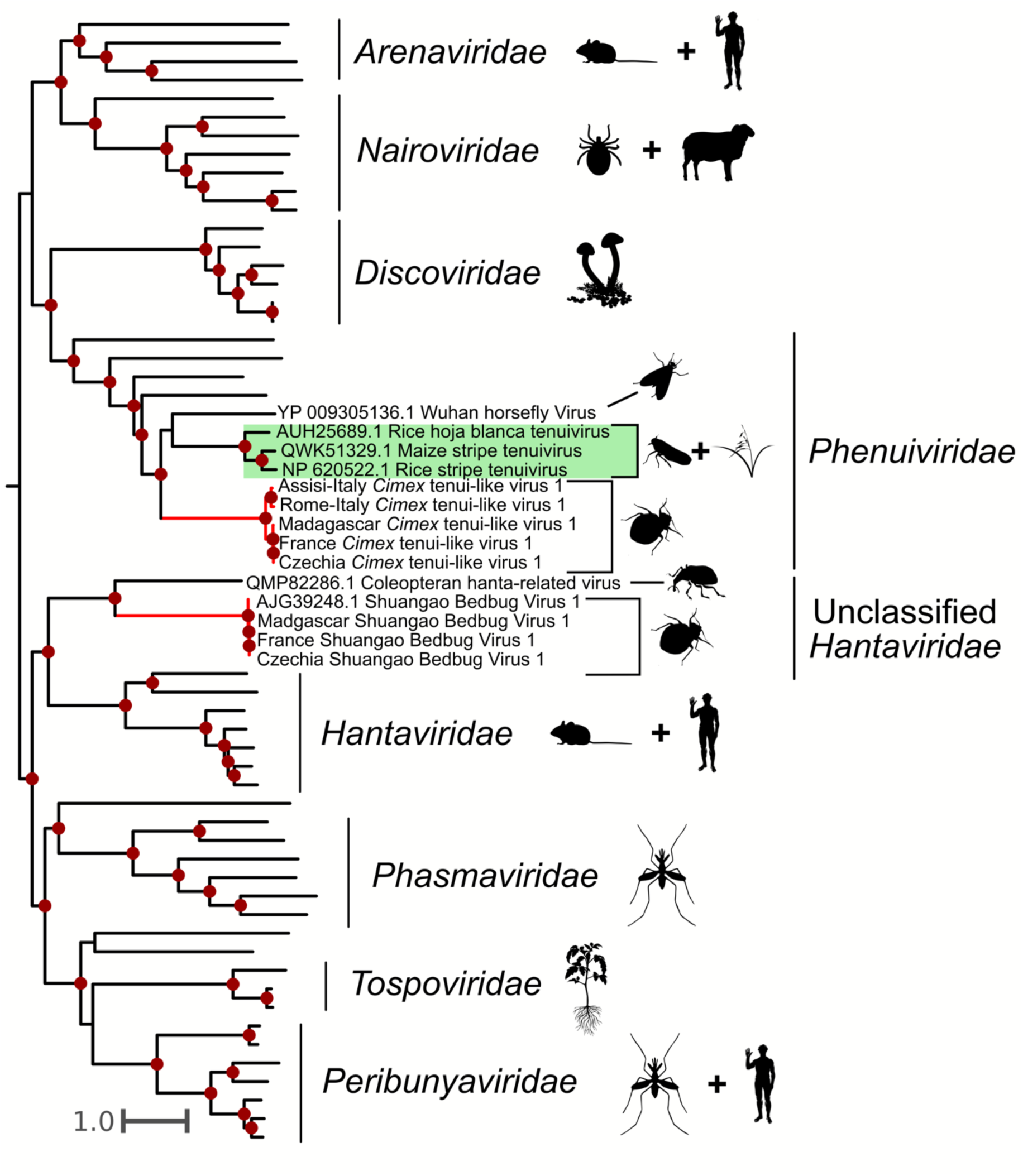
Phylogeny of the *Bunyavirales* including viruses detected in this study. Viruses found in this study are labeled by their location, and their branches are colored red. Dark red dots indicate bootstrap support greater than 75. Silhouette images represent the general host taxa of a virus or virus clade. The tree supports that we detected Shuangao bedbug virus 1 in multiple samples, and that it branches sister to the *Hantaviridae* (a clade of viruses that infects vertebrates) along with another insect-associated virus. *Cimex* tenui-like virus 1 groups within the family *Phenuivirudae*. Notably, *Cimex* tenui-like virus 1 branches sister to a clade including the *Tenuiviridae* genus (highlighted in green), which is comprised of important hemipteran-transmitted plant viruses.

### 3.3 Shuangao Bedbug Virus 2

Sbbv2 is described as an unclassified -ssRNA virus (NCBI:txid1608072) with a 10,925 bp monopartite genome. Our analyses show that it groups with the *Rhabdoviridae* family (**Figure 4**), consistent with the findings of Li *et al*. (2015). We detected Sbbv2 sequences in *C. hemipterus* samples from Madagascar samples 1 and 3. The genomes we assembled of this virus were around 3 kb longer than those from Li et al. (2015), ranging from 12,802-13,477 bp. Our results complete the genome of Sbbv2, as the extra 3 kb contains ORFs at the beginning of the genome: one encoding for a rhabdovirus nucleoprotein (CDD E-value = 1.77e^−06^), and the other encoding for what we suspect is the rhabdovirus phosphoprotein (**Supplementary Figure 1**). Although the latter ORF did not have any significant BLAST hits or conserved domains, we assume that it is the phosphoprotein based off synteny to other rhabdovirus genomes which generally encode for five proteins in the following order: nucleoprotein, phosphoprotein, matrix protein, glycoprotein, and large protein (Walker et al., 2022). Our study also extends the geographic range of this virus, as it was previously only detected in bed bugs from China (Li et al. 2015).

**Figure 4:**
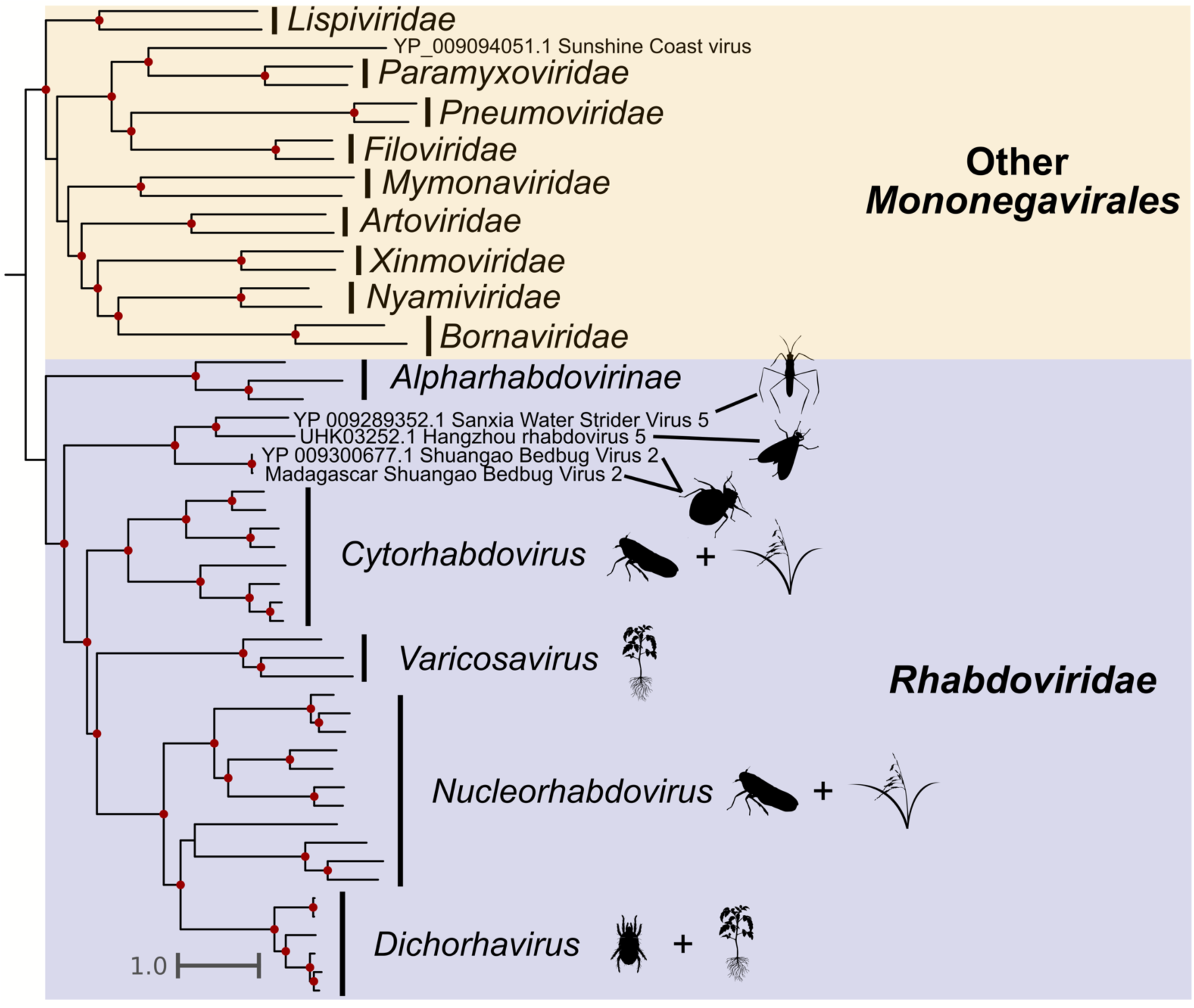
Phylogeny of the Mononegavirales with expansive *Rhabdoviridae* representation. Red dots indicate bootstrap support greater than 75. Sbbv2 is shown by a red branch. Non-rhabdovirus mononegaviruses are shown in yellow, while rhabdoviruses are shown in blue. Silhouette images represent the general host taxa of a virus or virus clade. Our phylogeny supports that Sbbv2 branches with other insect-associated viruses before the cytorhabdoviruses, nucleorhabdoviruses, and other plant infecting *Rhabdoviridae*.

### 3.4 Tenuiviridae

We detected sequences that had significant BLAST hits to viruses in the family *Phenuiviridae* (-ssRNA) in *C. lectularius* samples from Czechia, France, Rome-Italy, and Assisi-Italy, and *C. hemipterus* samples in Madagascar. These viruses are typically multisegmented, and from our BLASTx-based analysis, we detected a ∼9000 bp putative L segment. We found complete L segments in Czechia sample 3, Madagascar sample 2, Rome-Italy sample 1, and Assisi-Italy samples 1, 2, and 3. We found partial L segments in Czechia sample 2, France, and Rome-Italy sample 3. Our phylogenetic analysis groups these sequences in the family *Phenuiviridae,* sister to a clade including the tenuiviruses, which are a genus of hemipteran-transmitted plant infecting viruses (**Figure 3**). We refer to this putative virus as *Cimex* tenui-like virus 1.

Because bunyaviruses are typically multisegmented, we used an approach previously described in Walt et al. (2023) to identify candidate genomic segments based off sample co-occurrence. We found a single transcript that met our requirements with a *V_co_*= 0.86 and a *T_co_* = 0.75. The transcript is 2,851 nt long and encodes for a 902 amino acid protein. Based on transcript length in comparison to *Cimex* tenui-like virus 1’s closest BLAST hit (*Solenopsis invicta* virus 14: M segment length=2,705 bp), we propose this may be the M segment, which encodes a glycoprotein. InterProScan predicted two transmembrane domains in the amino acid sequence of this transcript, which is similar to the glycoprotein of *Solenopsis invicta* virus 14 as it also has two transmembrane domains. These results need further confirmation, as this transcript had no significant BLAST alignments to other known sequences, no predicted conserved domains in the translated protein, and no similar transcripts were found in the Italy samples, which also had *Cimex* tenui-like virus 1 sequences in them.

### 3.5 Luteoviridae

We detected a ∼2800 bp luteo-like virus 1 sequence (+ssRNA) in the Harlan lab strain, Czechia, and France samples, all of which were *C. lectularius*. We found complete transcripts in Czechia samples 1,2, and 3, France, and Harlan Strain-USA samples 1 and 2, and found a partial transcript in Harlan Strain-USA sample 3. Our phylogenetic analysis groups these viruses within a clade of unclassified *Luteoviridae*, sharing a common ancestor with 4 viruses detected in mosquitoes, and 1 virus detected in an anal swab from a bird (**Figure 5**). This clade branches sister to a group of three viruses, *Miscanthus* yellow fleck virus, Rabbit luteovirus, and *Arracacha* latent virus E. Rabbit luteovirus was discovered in a rabbit but was assumed to have come from its diet (Tsoleridis et al., 2019), and the other two viruses are plant viruses that are likely transmitted by aphids, another hemipteran insect (Bolus et al., 2020). We refer to this putative virus as *Cimex* luteo-like virus 1.

**Figure 5:**
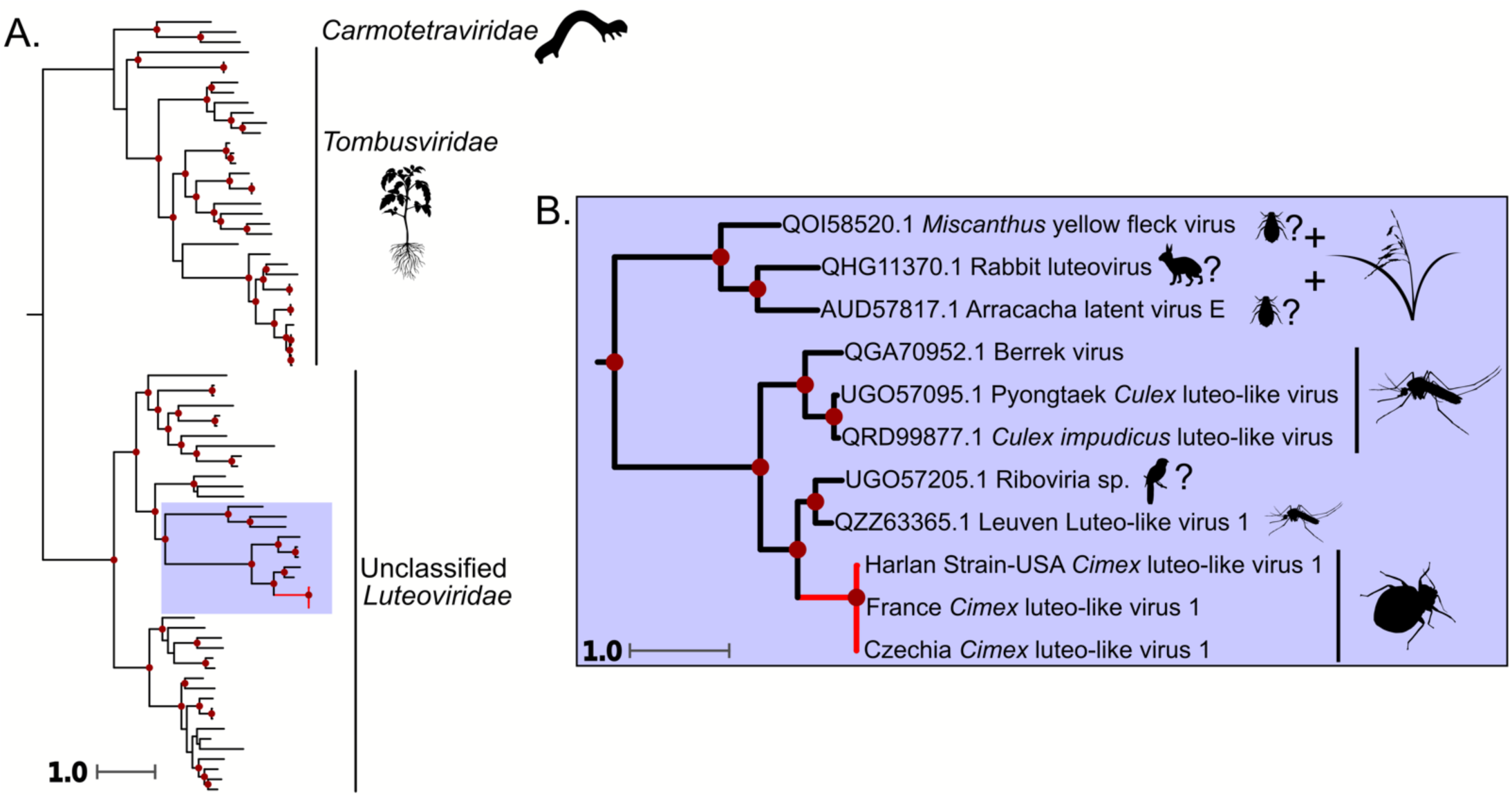
Phylogeny of the *Tolivirales*, including the ssRNA(+) virus discovered in this study. **A.)** Full phylogeny of the *Tolivirales*. The red branch indicates the *Cimex* luteo-like virus 1. Red dots along the tree show bootstrap support greater than or equal to 75. **B.)** Zoomed in view of the clade that *Cimex* luteo-like virus 1 groups in. Each virus is labeled by the respective location in which it was found. Dark red dots indicate bootstrap support greater than 75. Silhouette images represent the general host taxa of a virus or virus clade. *Cimex* luteo-like virus 1 groups with a clade of mosquito-associated viruses, sister to a clade of plant viruses that are putatively transmitted by aphids.

### 3.6 Totiviridae

We detected a toti-like virus (dsRNA) sequence in *C. lectularius* samples from five locations: the Harlan lab strain-USA, Czechia, France, Assisi-Italy, and the UK. The complete genome is around 7.8 kb and encodes for two large ORFs. One ORF (∼3880 nt) has significant BLASTx hits to other toti-like virus RdRps, and the other (∼1860 nt) has significant BLAST hits to other toti-like virus proline-alanine rich proteins (Spear et al., 2010). We recovered whole or nearly complete genomes for the UK (all samples), Czechia (complete for samples 1 and 3, partial for sample 2), France, and all three Assisi-Italy samples. Phylogenetically, this virus shares a most recent common ancestor with an unclassified totivirus isolated from a flea. The virus forms a clade with other Hemiptera and Thysanoptera associated toti-like viruses along with two viruses isolated from plants (**Figure 6**). We hereon refer to this putative virus as *Cimex* toti-like virus 1.

**Figure 6:**
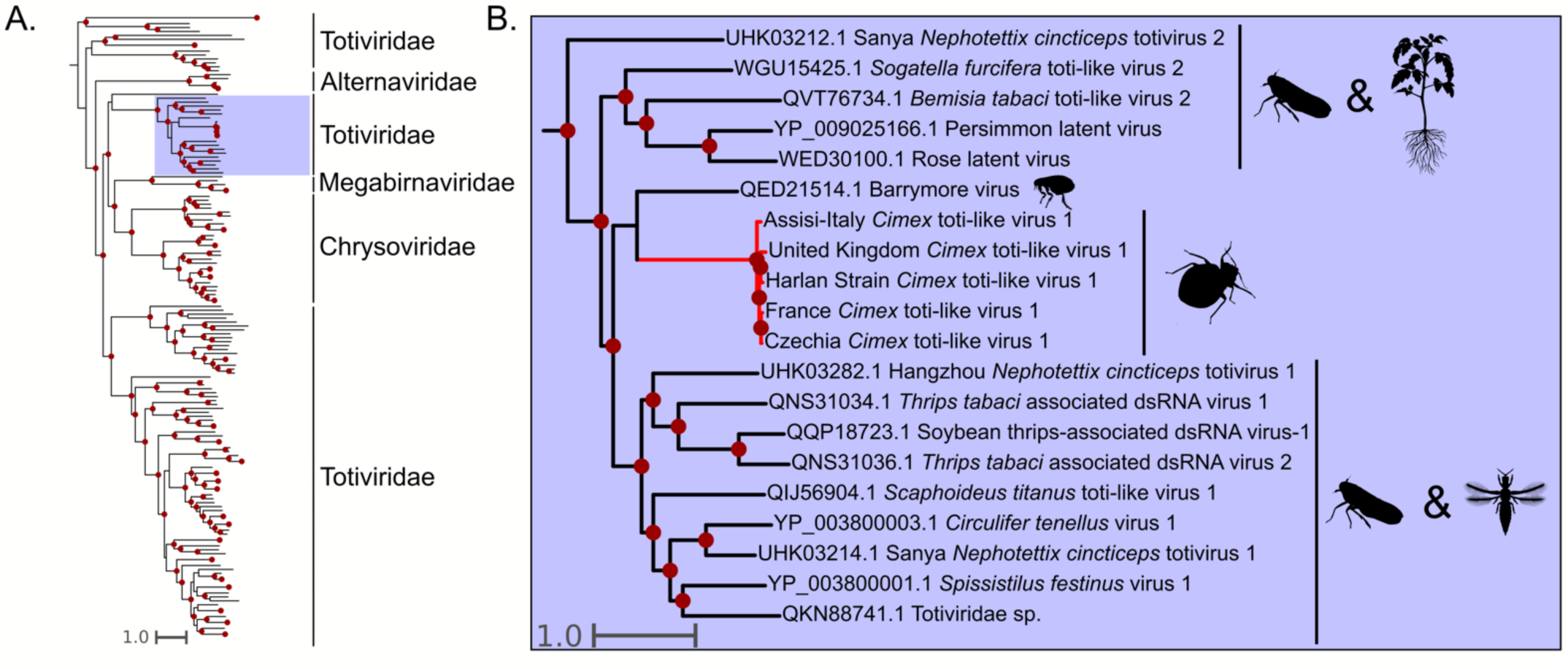
Phylogeny of the *Ghrabrivirales*, including the dsRNA virus discovered in this study. **A.)** Phylogeny of the *Ghrabrivirales* including diverse taxa from all families. *Cimex* toti-like virus 1 is shown on red branches and the clade it groups with is highlighted in red. Dots indicate a bootstrap value of 75 or above. **B.)** Zoomed-in view of the clade that *Cimex* toti-like virus 1 groups in. Silhouette images represent the general host taxa of a virus or virus clade.

### 3.7 Phylogeography of Bed Bug Viruses

All the virus sequences that we detected were distributed across geographically distant locations, so we investigated viral diversity between localities. We hypothesized that viral evolutionary relationships would reflect the differences in host species, and that within a host species, phylogenetic relationships would be reflective of geographic distance (Ballinger et al., 2022; Longdon et al., 2014). Specifically, we would expect *C. lectularius* and *C. hemipterus* to harbor closely related but distinct viral populations, and within *C. lectularius*, we would expect samples from Europe to be distinct from those rom North America. We only used sequences with complete RdRp domains and only included the unique sequences within individuals for our phylogeographic analysis.

#### 3.7.1 *Cimex* tenui-like virus 1

We found *Cimex* tenui-like virus 1 sequences in both *C. lectularius*, and *C. hemipterus*. In the case of *C. lectularius*, sequences from this virus were found in one individual from Rome-Italy, all three individuals from Assisi-Italy, one individual from Madagascar, the individual from France, and one individual from Czechia. Interestingly, the Italian samples form a distinct clade from the rest of the world (**Figure 7A**). When we grouped the samples by clade: Italy, world, and outgroup and assessed evolutionary distance between the groups, the mean p-distance between the Italian clade and the rest of the world clade was 22.4%, while the “within group” mean p-distance was 3.0% and 1.1% for the Italian clade and the rest of the world, respectively (**Table 1**). Interestingly, the “within group” distance is three times higher in the samples that come from the same country (Italy) than those collected in different countries (France, Chechia, and Madagascar).

**Figure 7:**
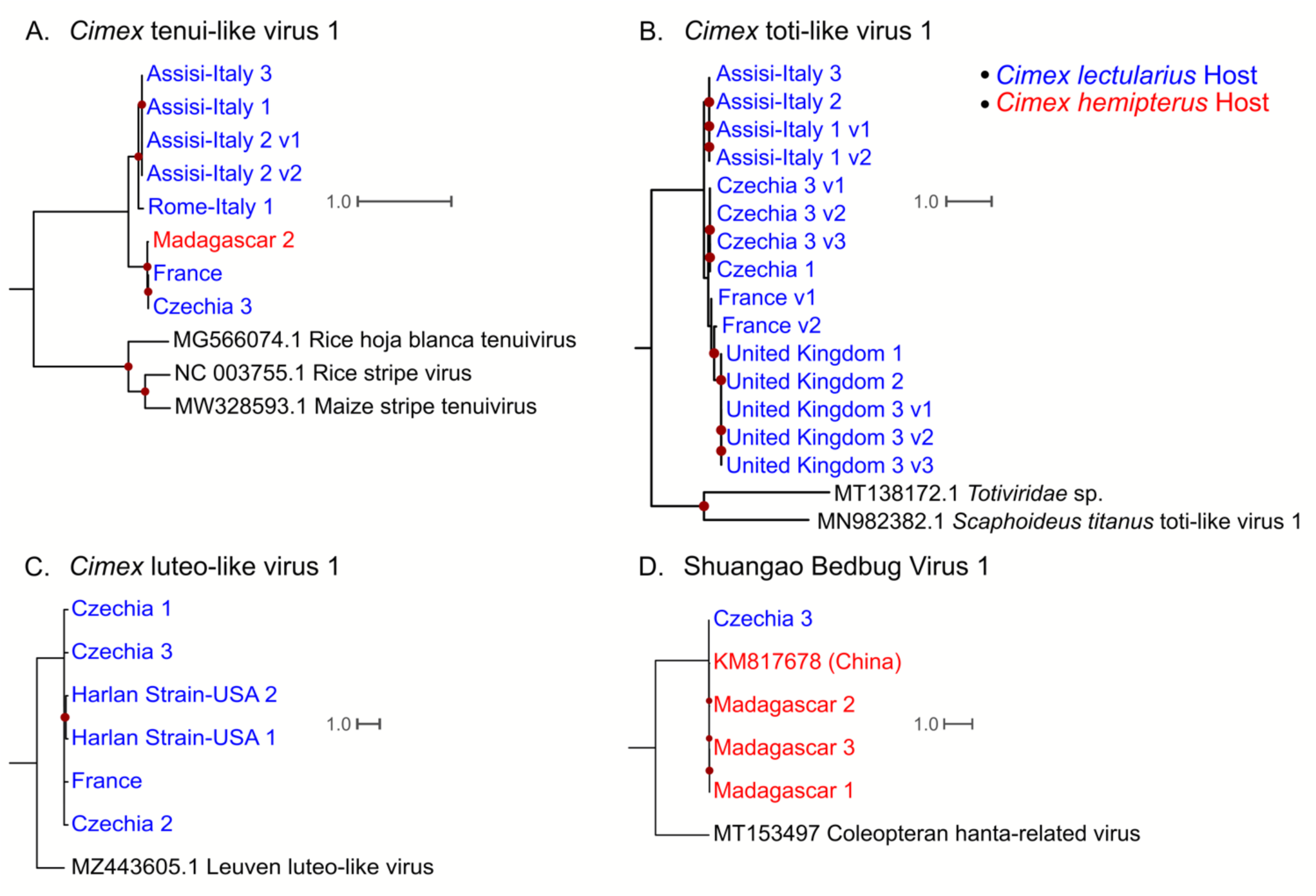
Phlyogeographic analysis of the putative bed bug viruses detected in this study. Only the coding sequences where an RdRp domain was detected were used. All trees were rooted using the closest phylogenetic neighbors from phylogenetic analyses in Figures 4-7. Dark red dots indicate bootstrap support greater than 75. Samples from *C. lectularius* hosts are indicated in blue text, and samples from *C. hemipterus* hosts are indicated in red text. **A.)** Phlyogeographic analysis of *Cimex* tenui-like virus 1. **B.)** Phylogeographic analysis of *Cimex* toti-like virus 1. **C.)** Phylogeographic analysis of *Cimex* luteo-like virus 1. **D.)** Phylogeographic analysis of Sbbv1.

**Table 1:** Mean evolutionary distance between distinct clades of viruses from phylogeographic analysis. Only *Cimex* tenui-like virus 1 and *Cimex* toti-like virus 1 are shown as they had highly supported trees containing many viral genomes. Distinct phylogenetic clades were formed from the Italy samples, the rest of the world, and the outgroups.

| *Cimex tenui*-like virus 1 |  | P-distance |  |  |
| --- | --- | --- | --- | --- |
|  |  | Italy | Rest of the World | Outgroup |
|  | Italy | 3.0% |  |  |
|  | Rest of the World | 22.4% | 1.1% |  |
|  | Outgroup | 55.8% | 55.9% | 36.5% |
| *Cimex toti*-like virus 1 |  |  |  |  |
|  | Italy | 0.3% |  |  |
|  | Rest of the World | 16.4% | 10.0% |  |
|  | Outgroup | 51.5% | 51.8% | 50.8% |

Because the samples collected in Italy had a higher “within group” mean than the samples from the rest of the world, we computed pairwise comparisons of p-distance for the samples used in the phylogeny in **Figure 7A** (**Table 2**). We found that one sequence of *Cimex* tenui-like virus 1 (Rome-Italy 1) is the driver of the viral genetic diversity within the group of samples collected in Italy, since the Assisi-Italy samples are identical, but the Rome-Italy 1 sample is 7.5% different from all the Assisi-Italy samples at the nucleotide level (**Table 2**). Furthermore, the France and Czechia 3 *Cimex* tenui-like virus 1 sequences are nearly identical, while the Madagascar 2 *Cimex* tenui-like virus 1 sequence is only 1.7% different from both the France and the Czechia samples (**Table 2**), even though the Madagascar samples are *C. hemipterus*, while the rest of the samples are *C. lectularius*.

**Table 2:** Pairwise p-distances of *Cimex* tenui-like virus 1 RDRP coding sequences. The matrix includes the outgroup sequences used in Figure 7A. RHBTV = Rice hoja blanca tenuivirus, RSV = Rice Stripe Virus, MSTV = Maize Stripe Tenuivirus.

|  | Rome-Italy 1 | Assisi-Italy 1 | Assisi-Italy 2 (1) | Assisi-Italy 2 (2) | Assisi-Italy 3 | Czechia 3 | France | Madagascar 2 | RHBTv | RSV | MSTV |
| --- | --- | --- | --- | --- | --- | --- | --- | --- | --- | --- | --- |
| Rome-Italy 1 | - |  |  |  |  |  |  |  |  |  |  |
| Assisi-Italy 1 | 7.5% | - |  |  |  |  |  |  |  |  |  |
| Assisi-Italy 2 (1) | 7.5% | 0.0% | - |  |  |  |  |  |  |  |  |
| Assisi-Italy 2 (2) | 7.5% | 0.0% | 0.0% | - |  |  |  |  |  |  |  |
| Assisi-Italy 3 | 7.5% | 0.0% | 0.0% | 0.0% | - |  |  |  |  |  |  |
| Czechia 3 | 23.1% | 20.6% | 22.6% | 21.3% | 22.6% | - |  |  |  |  |  |
| France | 23.1% | 22.6% | 22.6% | 20.6% | 22.6% | 0.0% | - |  |  |  |  |
| Madagascar 2 | 23.0% | 22.6% | 22.6% | 21.3% | 22.6% | 1.7% | 1.7% | - |  |  |  |
| RHBTv | 56.3% | 56.3% | 56.3% | 54.9% | 56.2% | 56.2% | 55.3% | 56.0% | - |  |  |
| RSV | 56.7% | 56.5% | 56.5% | 54.9% | 56.5% | 56.7% | 56.1% | 56.8% | 39.5% | - |  |
| MSTV | 55.1% | 55.4% | 55.4% | 54.4% | 55.4% | 55.4% | 54.8% | 55.4% | 39.3% | 30.7% | - |

#### 3.7.2 *Cimex* toti-like virus 1

We only found *Cimex* toti-like virus 1 sequences in *C. lectularius* samples. We found them in two of the three Czechia individuals, the France individual, all Assisi-Italy individuals, and all UK individuals. Once again, the samples collected from Italy form a distinct clade from the rest of the world (**Figure 7B**). We grouped the samples by the Italy clade, the rest of the world clade and the outgroup and found that there was little difference between the Italy viruses (“within group” mean p-distance = 0.3%), while rest of the world clade were more genetically distant (“within group” mean p-distance = 10%) (**Table 1**). The “between group” mean distance between the virus sequences detected in Italy and the virus sequences detected from the rest of the world is 16.4%. Within the rest of the world clade, the viruses from Czechia form one group and the France and UK viruses form another group, following expected geographic patterns.

#### 3.7.3 *Cimex* luteo-like virus 1

We only found *Cimex* luteo-like virus 1 sequences in *C. lectularius*, and we detected them in all three Czechia samples, two of the Harlan Strain-USA samples, and the France individual. The tree has low resolution other than one branch containing the viral sequences detected in the Harlan Strain-USA bugs (**Figure 7C**). All the *Cimex* luteo-like virus 1 RdRp-encoding transcripts have a mean p-distance of 4%.

#### 3.7.4 Shuangao Bedbug Virus 1

We detected Sbbv1 sequences in all three *C. hemipterus* individuals collected from Madagascar, and one *C. lectularius* individual from Czechia. The Madagascar samples and the previously described Sbbv1 sample (detected in China) group together with 100% support (**Figure 7D**). This reflects host taxonomy, as the Sbbv1 sample that was first detected in China was also detected in *C. hemipterus*. Because the bed bugs collected in Czechia were *C. lectularius,* our study expands the host range of Sbbv1 to *C. lectularius*. Overall, these viruses are very similar to each other with an average p-distance of 2% in the RdRp-encoding transcripts.

### 3.8 Influence of *Wolbachia* on Viral Abundance

Many hypotheses have been proposed to understand why bed bugs have never been linked to pathogen transmission, including one involving their *Wolbachia* endosymbiont as a potential factor (Pietri, 2020). *Wolbachia* has formed a nutritional symbiosis with bed bugs, as it provides them with B-vitamins (Hosokawa et al., 2010). Other studies have shown that Wolbachia colonization can confer resistance to viral infection in other insects such as *Drosophila*, mosquitoes, and even hemipteran insects (Cogni et al., 2021; Gong et al., 2020; Lindsey et al., 2018; Teixeira et al., 2008). A study investigating the influence of bed bug *Wolbachia* on feline calicivirus titer has been conducted, but there was no evidence that the virus ever replicated inside of the bed bugs (Fisher et al., 2019). To investigate the effects that *Wolbachia* may have on viral abundance, we mapped all reads to the viral genomes detected in this study and the *Wolbachia* endosymbiont of *Cimex lectularius* genome. **Table 3** shows the percentage of reads that mapped to the virus and *Wolbachia* genomes from each sample. We used these values to investigate the influence of Wolbachia reads on viral reads and we found no correlation (*y* = 0.41 - 0.042*x*, *R* = −0.14, *p* = 0.53) between percent *Wolbachia* reads and percent virus reads in a sample (**Figure 8**).

**Figure 8:**
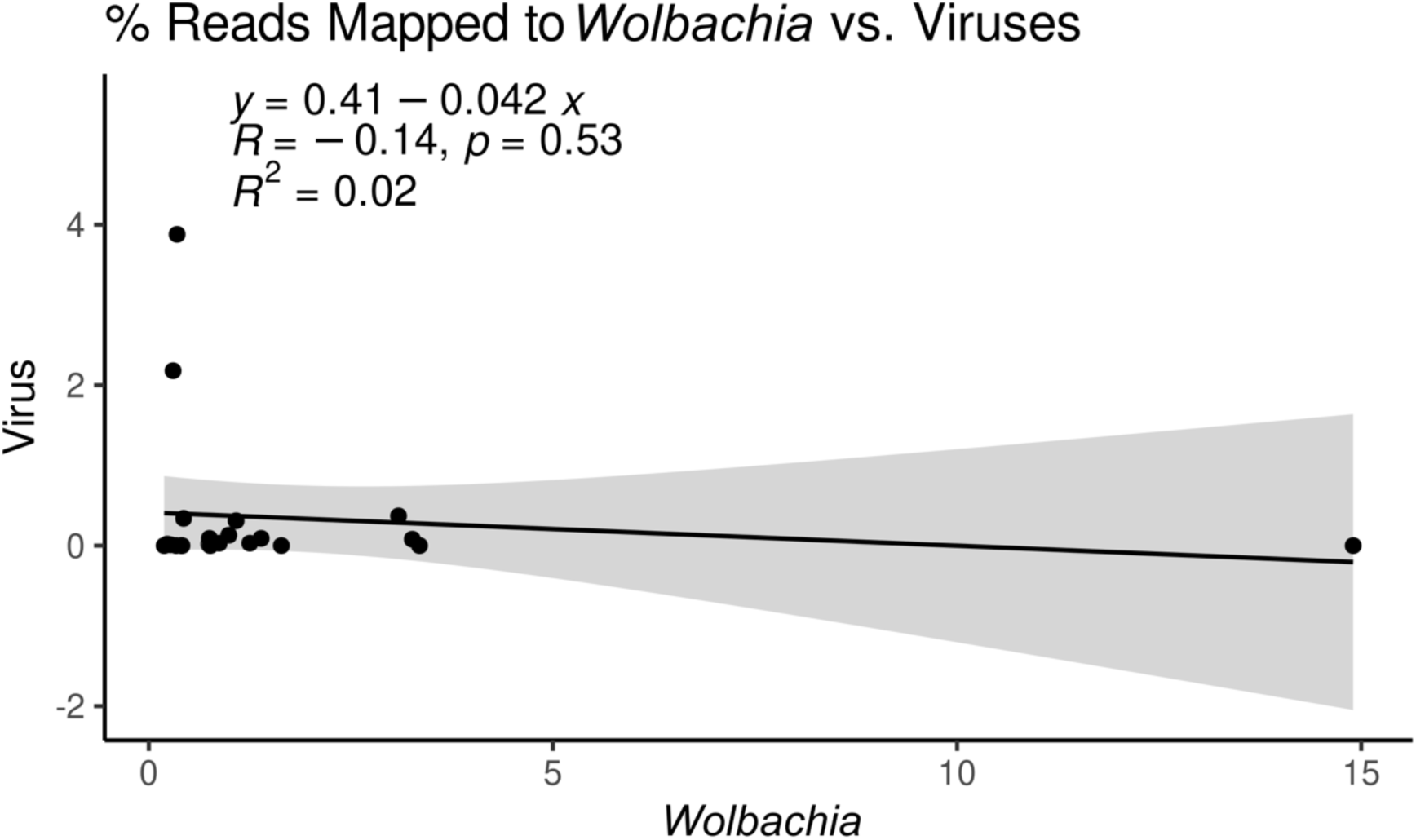
Correlation of the percentage of reads mapped to *Wolbachia* endosymbiont of *Cimex lectularius* and number of reads mapped to the viruses detected in this study. There is no correlation between percent of Wolbachia reads present in a sample and percent of RNA virus reads present in a sample.

**Table 3:** Percent reads mapped to the viruses detected in this study, or *Wolbachia* endosymbiont of *Cimex lectularius*.

| Sample | % Reads<br>Mapped to<br>Viruses | % Reads mapped<br>to *Wolbachia* |
| --- | --- | --- |
| Czechia 1 | 0.02 | 0.74 |
| Czechia 2 | 3.88 | 0.35 |
| Czechia 3 | 0.09 | 1.39 |
| Harlan Strain-USA 1 | 0.00 | 0.40 |
| Harlan Strain-USA 2 | 2.18 | 0.30 |
| Harlan Strain-USA 3 | 0.00 | 0.19 |
| France | 0.34 | 0.43 |
| Rome-Italy 1 | 0.01 | 0.76 |
| Rome-Italy 2 | 0.00 | 0.41 |
| Rome-Italy 3 | 0.00 | 0.34 |
| Assisi-Italy 1 | 0.01 | 0.27 |
| Assisi-Italy 2 | 0.09 | 0.75 |
| Assisi-Italy 3 | 0.02 | 0.23 |
| Madagascar 1 | 0.03 | 1.25 |
| Madagascar 2 | 0.03 | 0.87 |
| Madagascar 3 | 0.08 | 3.26 |
| Ohio-USA 1 | 0.00 | 14.90 |
| Ohio-USA 2 | 0.00 | 3.35 |
| Ohio-USA 3 | 0.00 | 1.64 |
| United Kingdom 1 | 0.37 | 3.09 |
| United Kingdom 2 | 0.31 | 1.08 |
| United Kingdom 3 | 0.13 | 0.99 |

## 4. Discussion

### 4.1 Detection and Phylogenetics of Bed Bug viruses

Bed bugs are a worldwide urban pest, that have undergone population resurgence for the last 20-30 years. Although their capacity to transmit human disease remains unknown, interest in their vector competence is high because of the increasing frequency in outbreaks (Doggett & Lee, 2023). We did not detect any known human viruses, but our study supports that metatranscriptomic surveillance is a useful technique to detect what known or emerging pathogens bed bugs could potentially transmit. We detected two previously known virus sequences associated with *C. hemipterus*, and 3 novel putative bed bug viruses.

The two previously detected bed bug viruses, Sbbv1 and Sbbv2 had been found in China associated with the tropical bed bug, *C. hemipterus* (C. X. Li et al., 2015). Our phylogeny groups Sbbv1 with an insect virus sister to the *Hantaviridae* (**Figure 3**). This agrees with the findings of Käfer et al. (2019) and supports the hypothesis that the *Hantaviridae* may have originated from arthropod viruses, and subsequently shifted to infecting vertebrate hosts (Marklewitz et al., 2015). Although Sbbv1 shares a common ancestor with the hantaviruses, it is unknown whether it is of concern to humans. We extend Sbbv1’s geographical range from China to Czechia, Madagascar, and France, and extend its host range to *C. lectularius*, as it had previously only been detected in the tropical bed bug, *C. hemipterus* (C. X. Li et al., 2015).

Sbbv2 provides an interesting insight to plant-infecting rhabdovirus evolution, as many economically important plant rhabdoviruses are transmitted by hemipteran insects (Whitfield et al., 2018). Our phylogeny agrees with Longdon et al. (2015), as Sbbv2 groups with insect-specific clade of rhabdoviruses that shares a common ancestor with the cytorhabdoviruses and the nucleorhabdoviruses (**Figure 4**). This supports the hypothesis that these viruses infected hemipteran insects before they infected plants (Longdon et al., 2015). It is interesting to note that Sbbv2 has persisted in the bed bug lineage despite shifts in feeding strategy from plants to other insects, to obligate blood feeders (Johnson et al., 2018). Our bunyavirus phylogeny (**Figure 3**) also suggests that the tenuiviruses, which are important insect-transmitted plant viruses, infected their insect hosts before evolving the ability to infect plants. This is indicated by *Cimex* tenui-like virus 1 branching sister to the tenuiviruses, and another insect virus, Wuhan horsefly virus, branching with this group (**Figure 3**).

Totiviruses are dsRNA viruses that typically infect fungi, but there is a growing number of toti-like viruses detected in arthropod and vertebrate metatranscriptomic studies (Tighe et al., 2022). According to our phylogenetic analysis, *Cimex* toti-like virus 1 falls within a clade of arthropod and plant infecting viruses, along with a toti-like virus 1 detected in an anal swab from a bird (GenBank: QKN88741.1) (**Figure 6**). Interestingly *Cimex* toti-like virus 1 shares a most recent common ancestor with a virus detected in fleas, which are also obligate blood feeders (Harvey et al., 2019). This supports the hypothesis that similarities in ecological niche could be more correlative of viral similarity than taxonomic relatedness (C. X. Li et al., 2015). Most other viruses in this clade are Hemiptera or Thysanoptera-associated (a sister group to the Hemiptera) viruses.

### 4.2 Phylogeography of bed bug viruses

Although our study design limited an extensive phylogeographic analysis, we found unprecedented patterns of viral diversity. First, we found that bedbug viruses detected in this study are not geographically restricted and can infect more than one host species. We detected sequences from four of the five viruses in this study intercontinentally and sequences from two out of five viruses were found in both *C. lectularius* and *C. hemipterus*. (**Figure 7**). Second, if we found viruses that were present in both bed bug species, we expected these viruses to form distinct clades reflecting bed bug taxonomy, as differences in host receptors between species would add selective pressures to viral infection (Longdon et al., 2014). Our results did not match that expectation, as the *Cimex* tenui-like virus 1 detected in the *C. hemipterus* (Madagascar) samples grouped with the *Cimex* tenui-like virus 1 sequences from *C. lectularius* (France and Czechia), while other *Cimex* tenui-like virus 1 sequences detected in *C. lectularius* formed their own distinct clade (Rome-Italy and Assisi-Italy) (**Figure 7A**). Third, we expected that viruses from similar geographic location would form distinct phylogenetic groups. This trend was generally followed, but strikingly, in every case where viral sequences were detected in samples from Italy, the sequences form their own distinct clades separated from the rest of Europe (**Figure 7A&B**). Furthermore, even though the *Cimex* tenui-like virus 1 sequences from Italy group together phylogenetically, there is higher mean evolutionary distance within these samples than within the *Cimex* tenui-like virus 1 sequences from the rest of the world, which include samples from France, Czechia and Madagascar, and include two different host species (**Tables 1 & 2**). Previous studies of bed bug phylogeography have found low genetic diversity within bed bug infestation sites, but high genetic diversity between infestation sites even of relatively close proximity, which could be due to their dependence on humans for dispersal (Fountain et al., 2014; Saenz et al., 2012). Bed bug dispersal by humans could also explain the unexpected patterns of bed bug virus phylogeography, as the distinct groups of bed bug viruses could be explained by Italy’s popularity as a travel destination, with Rome being a hotspot for tourism, and Assisi being a frequent site of pilgrimage. Along with this, the similarity between viruses detected in Madagascar and Europe could reflect travel between these two places.

### 4.3 *Wolbachia* influence on viral abundance

As an additional exploration of our dataset, we investigated if the amount of *Wolbachia* reads was correlated with viral abundance. It has been hypothesized that *Wolbachia* could have a protective effect against viral infection in bed bugs, similar to what has been seen in mosquitoes and *Drosophila* (Cogni et al., 2021; Fisher et al., 2019; Hussain et al., 2023; Lindsey et al., 2018; Teixeira et al., 2008; Terradas & McGraw, 2017). We used percentage of reads mapped to *Wolbachia* and percent mapped to the viral genomes detected in this study as a proxy of abundance. We found that there was no correlation between *Wolbachia* and virus abundance when a potential outlier sample was present (**Figure 8**). These results indicate that unlike Diptera-associated *Wolbachia,* bed bug *Wolbachia* may not confer viral resistance. Although our experimental design was not ideal to test *Wolbachia’s* influence on bed bug virus fitness, these results provide a preliminary look into how *Wolbachia* may affect viruses that replicate inside of bed bugs.

## 5. Conclusions

Our study opens interesting questions about the bed bug virosphere but does not provide evidence that bed bugs transmit human viruses. On the contrary, humans may drive bed bug virus diversity by facilitating dispersal and local extinction of host populations (Fountain et al., 2014). Future studies should assess the pathogenicity and transmission routes of these viruses to have a more comprehensive understanding of their potential in biocontrol or as emerging diseases. Along with this, a more comprehensive sampling strategy and phylogeographic analysis could shed light on the interesting patterns of bed bug virus dispersal.

## ACKNOWLEDGMENTS

The authors would like to thank Matthew J. Ballinger for his helpful insight and comments on our manuscript.

## DATA AVAILABILITY

All newly reported virus genome sequences used in this study will be made available in GenBank, and all reads generated in this study will be deposited to NCBI’s Sequence Read Archive (SRA) database.

## SUPPLEMENTARY DATA

Supplementary data will be available online.

