## Supplemental Figure 1 for "Do bed bugs transmit human viruses, or do humans transmit bed bug viruses? A worldwide survey of the bed bug RNA virosphere" (from another version of this preprint: version 2, 2023-10-28)

**SUPPLEMENTARY DATA:**

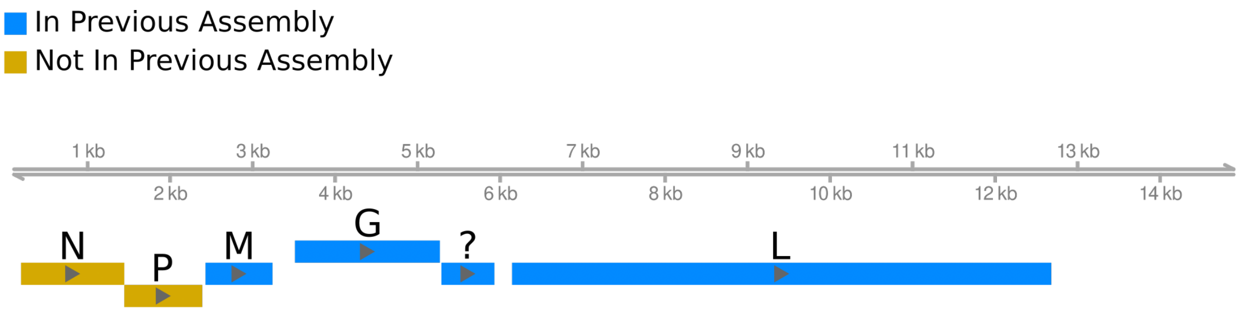


**Supplementary Figure 1: Completion of Shuangao bedbug virus 2 (Sbbv2) genome.** The previous assembly of Sbbv2 (Refseq: GCF_001755405.1) was missing the nucleoprotein and phosphoprotein typically found in rhabdovirus genomes (shown in blue). Our assembly was around 3 kb longer than the original assembly and included two extra ORFs predicted to be the nucleoprotein and the phosphoprotein (shown in yellow).
